# The Safety of a Therapeutic Product Composed of a Combination of Stem Cell Released Molecules from Adipose Mesenchymal Stem Cells and Fibroblasts

**DOI:** 10.1101/2020.02.14.950055

**Authors:** Greg Maguire, Peter Friedman

## Abstract

Stem cell transplants have demonstrated life-saving capabilities for some blood diseases, and the molecules and exosomes released from stem cells are currently in therapeutic development for a number of diseases and conditions, including neurodegenerative diseases, heart conditions, glaucoma, hearing loss, and skin diseases. Stem cell science is a relatively new science, and therapeutic development using stem cells, even approved stem cell therapies for blood diseases, is in need of a better understanding of mechanisms of action and acute and long-term safety profiles. Here we performed a number of safety tests for a stem cell-based therapeutic comprised of the stem cell released molecules from a combination of adipose derived mesenchymal stem cells and fibroblasts that have demonstrated efficacy in a number of conditions. Using in vitro, in vivo, and skin sensitivity studies in humans, the stem cell therapeutic comprised of stem cell released molecules was shown to have an excellent safety profile when tested for toxicity, mutagenicity, tumorigenesis, ocular toxicity, inflammation, and irritation.

## Introduction

Although stem cell transplants have been used for over 40 years with demonstrated life-saving capabilities for some blood diseases (Thomas et al, 1959), and the molecules and exosomes released from stem cells are currently in therapeutic development for a number of diseases and conditions, including neurodegenerative diseases (Maguire et al, 2019), heart conditions (Marban, 2018), glaucoma (Klingborn et al, 2017), hearing loss (Qui and Qui, 2019), and skin diseases (Wang et al, 2017; Maguire et al, 2019), stem cell science is a relatively new science, and therapeutic development using stem cells, even approved stem cell therapies for blood diseases, is in need of a better understanding of mechanisms of action and acute and long-term safety profiles, both for the cells and their released molecules. Approved bone marrow stem cell (BMSC) transplants have many associated risks, including possible induction of cancer (Cooley et al, 2000), including skin cancer (Omland et al, 2016), and aging of the tissue in which the implant occurs (Wood et al, 2016). Many factors, often overlooked, must be considered when developing stem cell-based therapeutics, including something as fundamental as the choice of stem cell type where adipose mesenchymal stem cells have many advantages over bone marrow stem cells for therapeutic development (Maguire, 2019). Here we performed a number of safety tests for a stem cell-based therapeutic comprised of the stem cell released molecules (secretome) from a combination of adipose derived mesenchymal stem cells (ADSCs) and fibroblasts (FBs), the combination of which has demonstrated efficacy in a number of conditions (Maguire et al, 2019; Maguire et al, 2019b), and is conceptually based on developing a systems therapeutic (Maguire, 2014) for the physiological renormalization of tissue in various disease states or abnormal conditions (Maguire et al, 2019). Using in vitro, in vivo, and skin sensitivity studies in humans, the stem cell therapeutic comprised of stem cell released molecules from ADSCs and FBs was shown to have an excellent safety profile when tested for in vitro and in vivo toxicity, the Ames mutagenicity assay, in vivo tumorigenesis, in vivo inflammation, ocular histology, and a human skin patch test for irritation and allergic reaction.

## Methods

Stem cell culture procedures have been described previously (Maguire et al, 2019). Briefly, a proprietary collection of stem cell lines derived from human skin were cultured using no penicillin/streptomycin under hypoxic conditions. When cultures reached confluence, they were passaged for a limited number of times before disuse. Total conditioned medium from the multiple cell types, containing a soluble fraction and an exosome fraction, was harvested at each passage and the passages combined into one batch for product development. Parts of our stem cell technology used here are covered by US patents: 9545370; 9446075; 20140205563; 20130302273.

### Safety Studies, HRIPT and Canine RIPT

The Human Repeat Insult Patch Test (McNamee et al, 2008) was used to assess primary and accumulative irritation, and/or allergic contact sensitization, in human subjects, male and female, between the ages of 18 and 66. All subjects were free of skin disease, and were prohibited from using topical or systemic antihistamines and steroids beginning seven days prior to the onset and throughout the duration of the study. Measurements were performed by a trained rater technician. For the induction phase of the study, the patches. 0.1 g of the combinatorial secretome per 1X1 inch square Webril dressing, were applied 3 times weekly for 3 weeks for a total of 9 applications. The patches were removed 48 hours after application and evaluated before the a fresh patch was reapplied to the same area. For the challenge phase of the study, two weeks after the final induction phase patch was applied, an adjacent, virgin area of skin then received a fresh patch and was evaluated at 24 and 72 hours following application of the patch. Ratings were scored using the following table:

0- No visible skin reaction
1- Barely perceptible or spotty erythema
2- Mild erythema covering most of the test site
3- Moderate erythema, possible mild edema
4- Marked erythema, possible edema
5- Severe erythema, possible edema, vesiculation, bullae, and/or ulceration

For the canine RIPT study, 27 qualified subjects absent of skin disease or irritation and no use of medications, male and female, ranging from age 2 to 7 years old were studied. S2RM was applied to the belly of the dog, held for two minutes to allow the S2RM to absorb into the skin, and no patch applied. The rater was a practicing veterinarian, who used the same rater scale that was used in the HIRPT. Photos were taken following each application that was done 3 times weekly for three weeks. All 27 dogs finished the study.

### Oral Toxicity Studies

This toxicity study used 3 groups of 3 male and 3 female Sprague-Dawley rats (18 total). Once-daily oral (gavage) administration was as follows: Group 1 was administered the vehicle only once daily at 4.4 mL/kg/dose for 28 consecutive days. Group 2 was administered S2RM once daily at 0.44 mL/kg/dose for 28 consecutive days. Group 3 was administered S2RM once daily at 4.4 mL/kg/dose for 28 consecutive days. As a biomarker for a possible inflammatory response to the S2RM administration, all rats were subjected to blood collection (Red-Top) for ELISA testing of the pro-inflammatory cytokines IL-10, and IL-31, measured prior to the first dose (Day 1) and on the day following the last dose (Day 29).

## Results

### Batch Reproducibility – Total protein

The method of Bradford was used in all protein determinations (Ernst and Zor, 2010). The Bradford protein assay is a spectroscopic analytical procedure used to measure the concentration of protein in a solution. Samples, performed in duplicate, from a total of 15 batches of S2RM were analyzed. The variation between duplicates in all cases was less than 10%. Five samples of different S2RM batches, and 5 samples of the individual batches of SRM from the individual cell lines that make-up the S2RM, were analyzed for total protein. All samples were taken from frozen aliquots of batches previously used to make skin care products with demonstrated efficacy. All batches produced at least 500 ug/ml of total protein, with a high value in one batch of 630 ug/ml. The mean value of all batches was 554 ug/ml. The variability in all the batches was 20% or less. With the exception of one batch displaying a high protein count (630ug/ml), the rest of the batches had a variability of 12% or less.

### Skin Safety Testing, Irritation

Ninety-one (91) qualified subjects, male and female, ranging in age from 18 to 66 years, were selected for this evaluation. Fifty (50) subjects completed the study. The remaining subjects discontinued their participation for various reasons, none of which were related to the application of the test material. Of the 50 subjects completing the study, all were rated at 0 during both the induction phase and the challenge phase, indicating that the S2RM induced no immediate or long term irritation, or allergic reaction. Because the S2RM is also intended for therapeutic development to treat companion animals, we tested for cross-species irritation in canines. Similar to that for humans, no irritation or allergic reaction was observed in the 27 canines as all scored “0” throughout the testing period.

### Oral Administration Toxicity, Tumorigenesis, and Inflammation Results

No significant differences in organ weights were found in the male or female rats when compared to controls (Figure 1A and B). When compared to the control group, there were no biologically relevant changes from Day 1 to Day 29 for group average IL-10 or IL-31 in either SRM-treated group, suggesting that the S2RM did not induce an inflammatory reaction when administered orally. All tissues were also analyzed for tumor formation, and no tumors were found in any of the analyzed tissues (analyzed tissues are depicted in Figure 1) in any of the animals. As a general indicator of health, body weights were recorded prior to the first dose and weekly thereafter including the day of euthanasia (Figure 1C). When compared to the control group, there was mildly decreased body weight gain in high-dose (SRM at 4.4 mL/kg/dose) males and females over the course of the study. Body weight in the SRM low-dose group (Group 2, 0.44 mL/kg/dose) was comparable to the control group throughout the study in males and females. Therefore the general health and the appetite of the experimental animals was normal.

**Figure 1.**
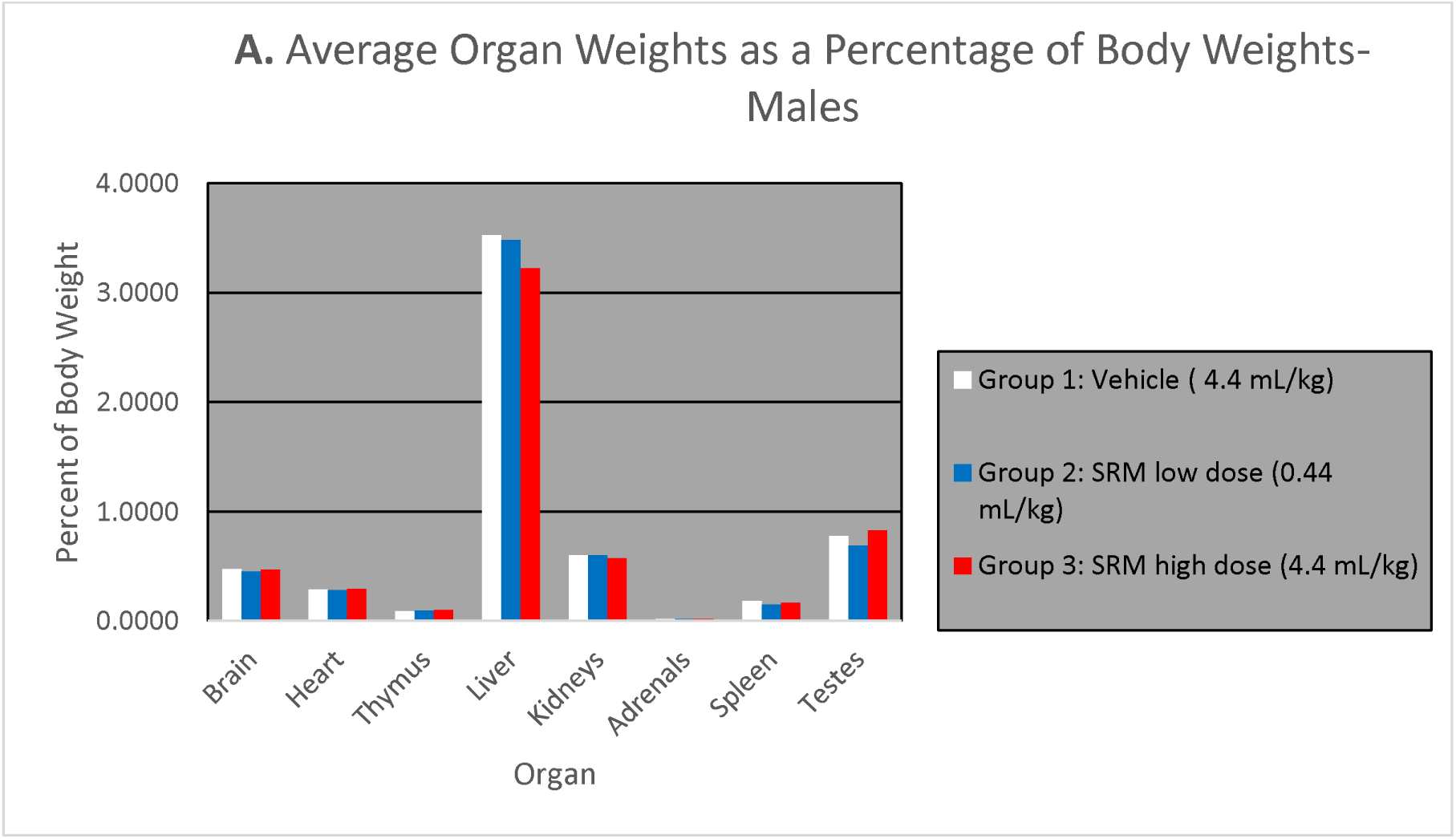
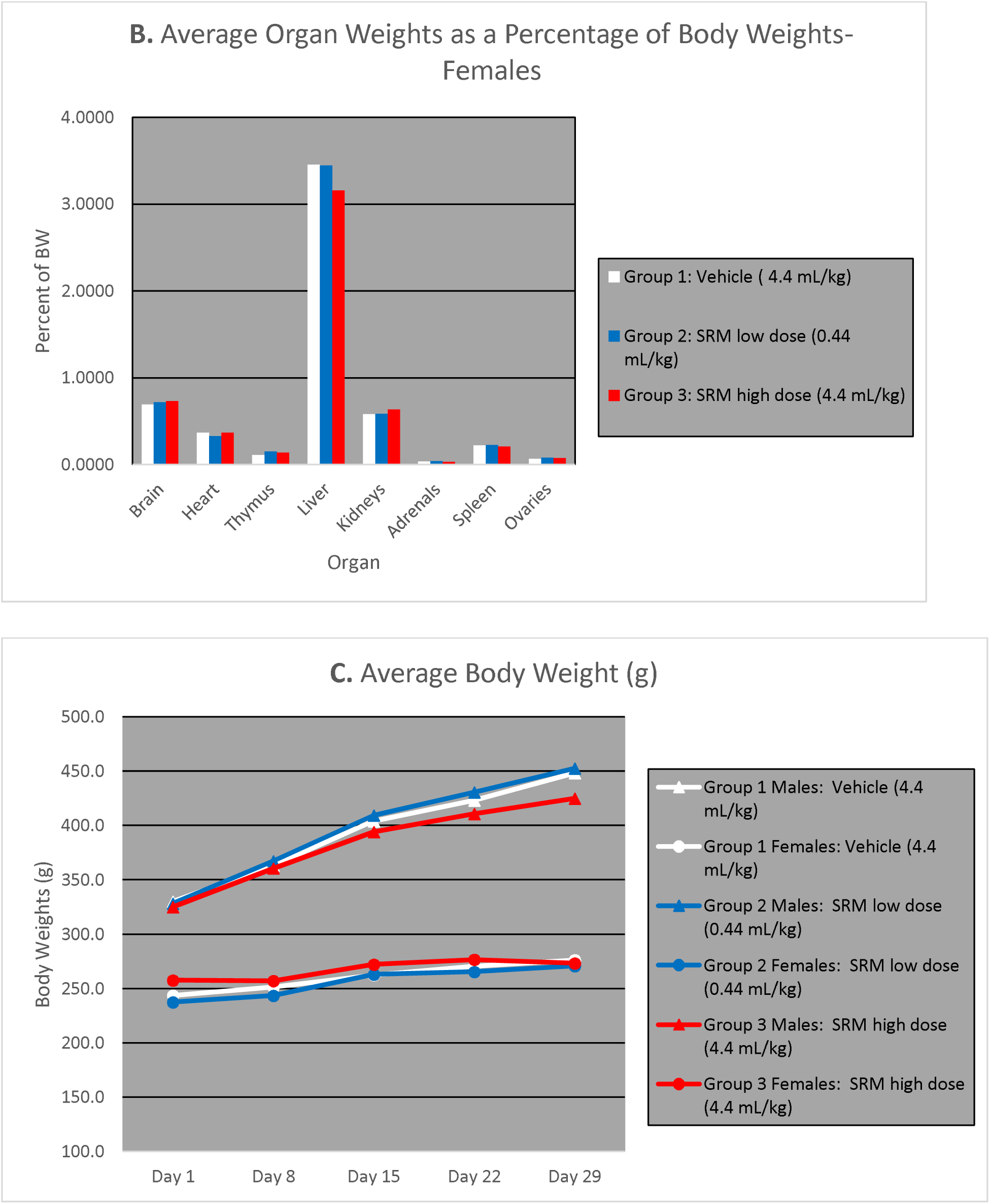
Average organ weights as a percentage of body weight for males, and B is that for females.C. Total body weight over a 30 day period during dosing by oral gavage of S2RM. No significant differences weights were observed for any of the organs, and none of the organs showed evidence of toxicity or tumorigenesis.

### Analysis of Ocular Pathology and Tumorigeneicity Following Topical Application of S2RM to the Eye

Both eyes were recovered from all animals following seven consecutive days of twice daily administration of the S2RM as eye drops applied to the cornea. Eyes were submitted to Colorado Histo-Prep for histopathological evaluation by a board-certified veterinary pathologist who had no knowledge of which eyes had been treated. Six rabbits (New Zealand White) received the S2RM solution in one eye, and the other eye served as a control. One control rabbit receiving no S2RM in either eye was also was also used. The samples were trimmed, processed, embedded, sectioned, and stained. Histopathology of the tissues was conducted on slides stained with hematoxylin and eosin with an emphasis on eye irritation and inflammation. No S2RM related lesions or tumors were observed in this study after 7 days of topical S2RM/Placebo treatment and prior to euthanasia. The eye globe sections were of very good quality and no differences in cornea morphology was detected in any specimen.

### Ames Test

The Ames reverse mutation was used in four strains of *Salmonella*. Mutagens identified using the Ames test are usually carcinogens, given Ames showed that 90% of carcinogens are also mutagenic (McCann et al, 1975). In the four strains of bacteria tested (Figure 1), the S2RM was shown to be non-mutagenic tested with S9 activation or without s9 activation.

**Figure 1.**
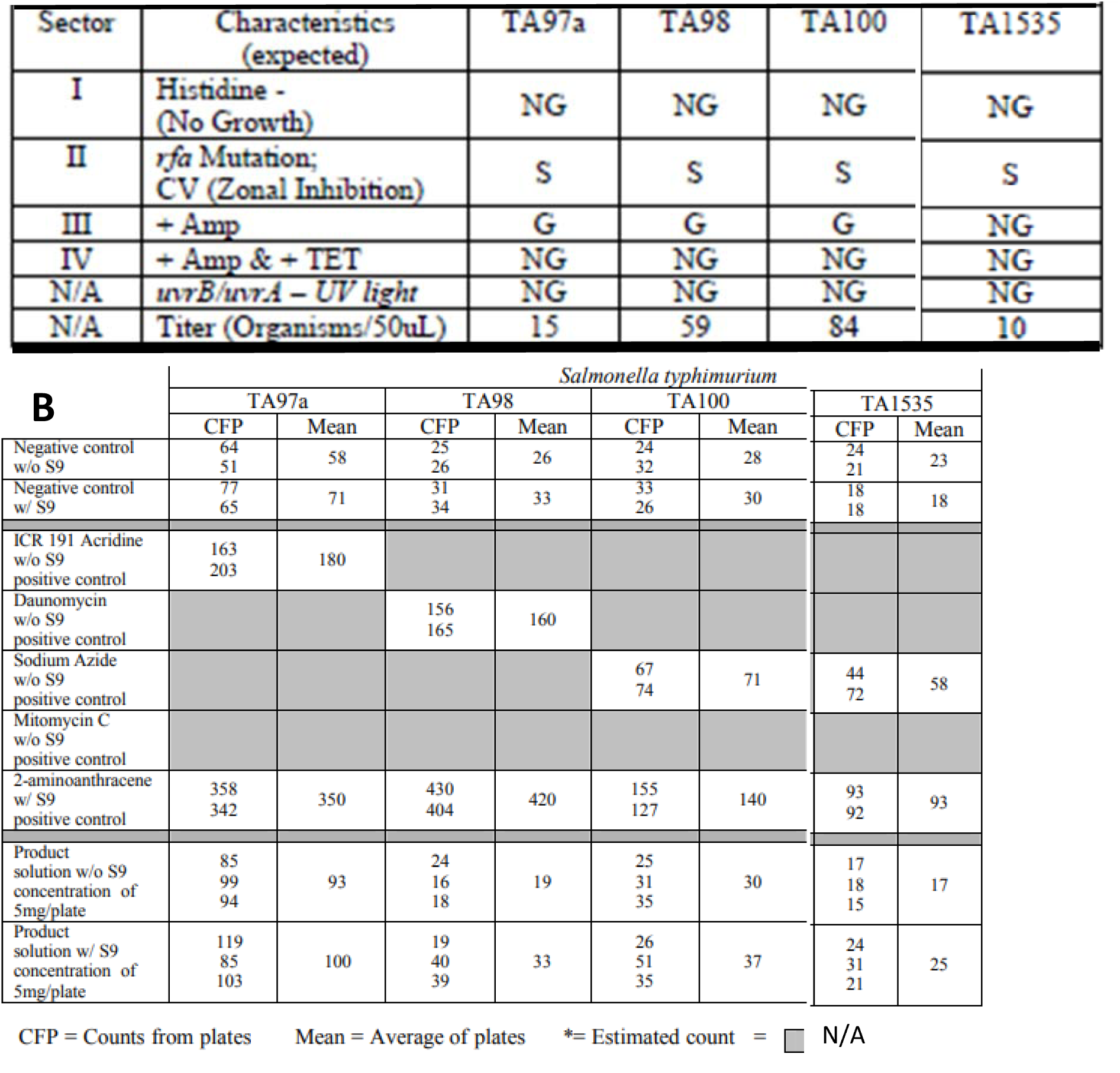
Ames test for mutagenicity in four strains of bacteria show that the S2RM is not a mutagen. NG = no growth, S= slight growth, G= growth, S= sensitive. A twofold or greater increase in the number of mean revertants for the S2RM over the mean number of revertants obtained from the negative controls was considered mutagenic. **A**. Strain characteristics and standard strain plate counts. **B**. Standard plate incorporation assay – reversion rates for tester strains.

### Cellular Cytotoxicity in a Mouse L929 Fibroblast Cell Line

Fibroblasts exists in most tissue compartments throughout the body, and particularly in the skin where the current S2RM technology has proven efficacious for a number of skin conditions (Maguire et al, 2019b). We therefore tested the S2RM for potential cytotoxic effects in fibroblasts. Table 2 shows the data providing evidence that the S2RM has no cytotoxic effects in fibroblasts as measured by observing changes in cell structure or the number of cells in the culture.

**Table 2.**
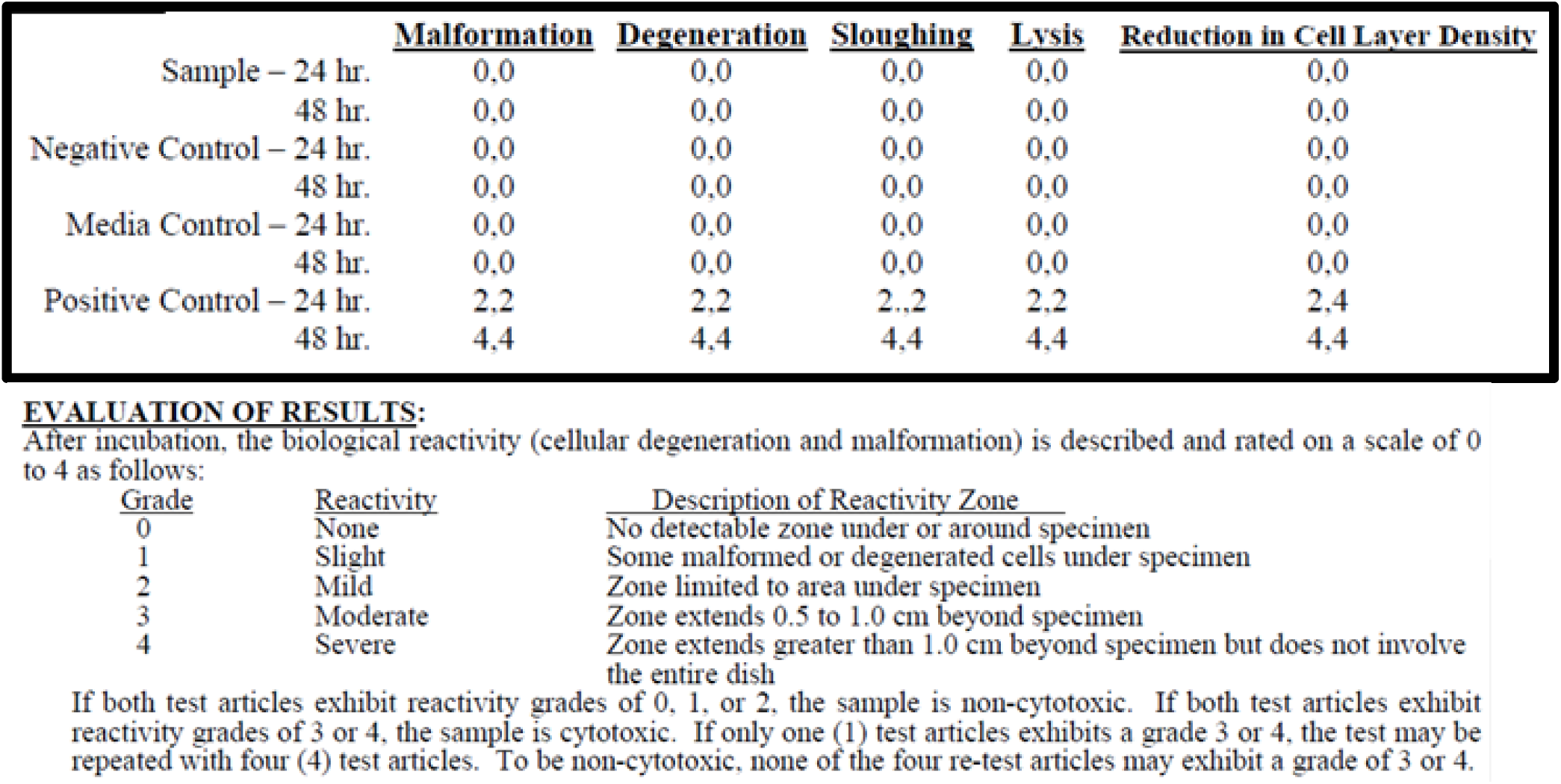
Using a mouse cell line L929, the S2RM conditioned media is shown to have no direct cytotoxic effects on the cells (fibroblasts).

**Table 3.**
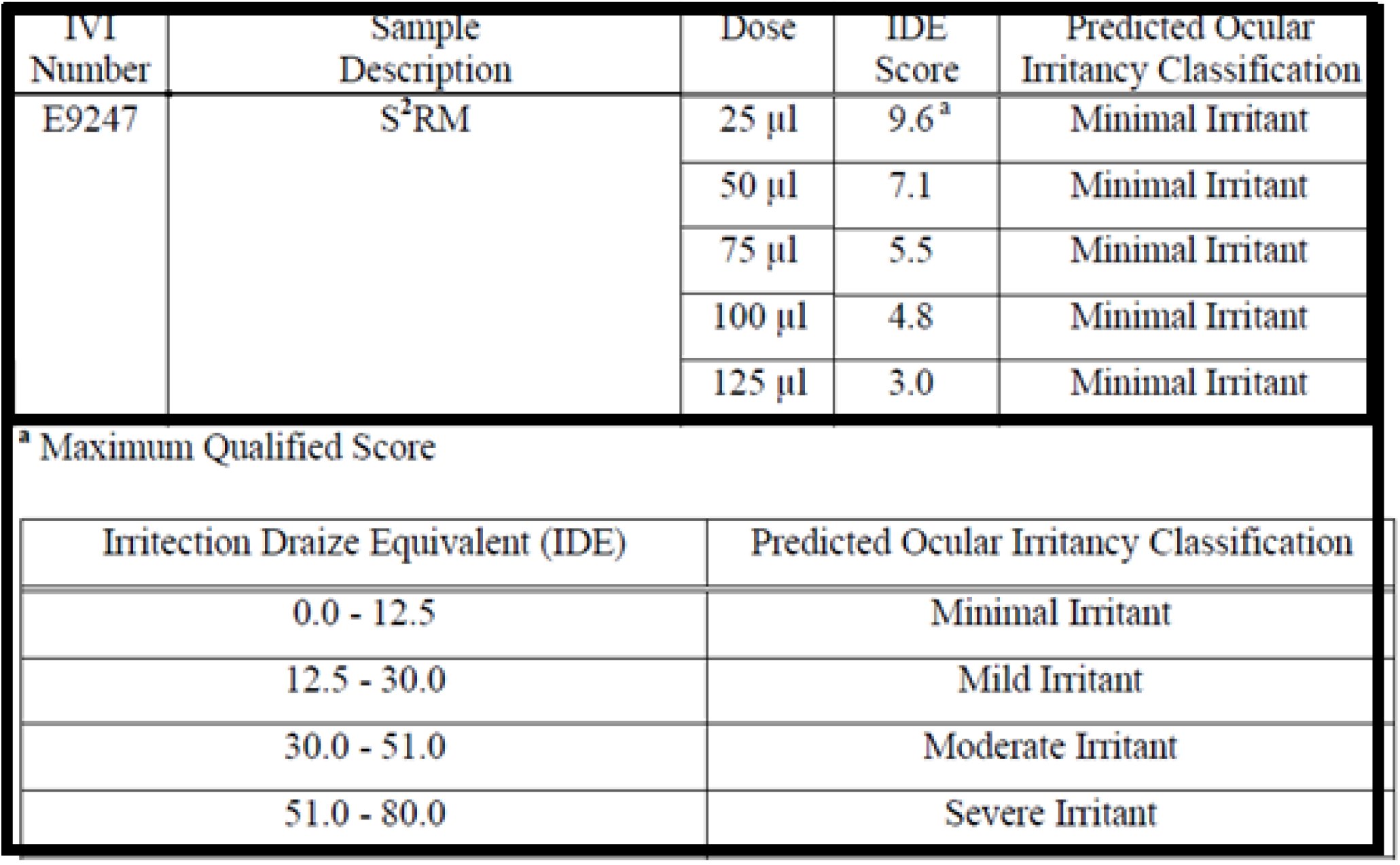
S2RM did not induce any misfolding or structural changes in the collagen, keratin, and other dermal proteins.

**Table 4.** Testing for the presence of four common heavy metals. None were present in significant, unsafe amounts.

| <u>Test</u> | <u>BTS Method No.</u> | <u>Result</u> |
| --- | --- | --- |
| Arsenic | M766.R04 | < 0.025 ppm* |
| Cadmium | M766.R04 | < 0.025 ppm |
| Lead | M766.R04 | < 0.025 ppm |
| Mercury | M766.R04 | < 0.025 ppm |

### Protein Misfolding Tests (Dermal Irritection® Assays System)

In this study the Irritection system from In Vitro International (Placentia, CA) was used for detection of misfolding and structural changes in proteins. Using this methodology, an irritant chemical will disrupt the ordered structure of keratin and collagen and result in the release of a bound indicator dye. Additionally, irritants will induce changes in conformation of the globular proteins found in the reagent solution. The extent of dye release and protein denaturation was quantitated by measuring the changes in optical density of the S2RM solution at 450 nm (OD450). The results provide evidence that S2RM induced no structural changes or misfolding of the test proteins.

### Heavy Metals

Numerous products and procedures are involved in processing the stem cells for collection of their molecules, presenting a number of points were contamination may take place. We therefor tested for the presence of four common heavy metals in the S2RM end product used as the therapeutic. No significant level of heavy metals was present in the S2RM.

## Discussion

The results presented her for a combination therapeutic containing the stem cell released molecules from ADSCs and FBs provide evidence that these molecules are safe whether they are administered orally, or topically to the eye or skin. No signs of toxicity, tumorigenesis, mutagenicity, irritation and allergic reaction, or inflammation were measured when the S2RM was compared to controls. These data are also important for stem cell transplants because when transplanted stem cells graft in the tissue and remain alive and healthy for a long time (Scala et al, 2018), much of the cell’s therapeutic effect is mediated by the release of molecules that act as paracrine and autocrine factors (Chimenti et al, 2010; Riazifar et al, 2017). Understanding the cellular effects versus the paracrine effects of stem cell therapy is important for the development of therapeutics, both in terms of efficacy as well as safety. For example, we know that bone marrow stem cell transplants induce aging of the implanted tissue (Wood et al, 2016) and increase the probability of cancer (Cooley, 2000; Christopher, 2018), but we don’t understand the mechanisms by which this happens. Is it the choice of the bone marrow stem cell (BMSC) type versus the adipose mesenchymal stem cell (ADSC) type that is responsible for these issues (Maguire, 2019), or the implantation process of exogenous cells, or the release of specific molecules from the BMSCs that cause these adverse events? While we have no evidence to answer the aforementioned questions directly in this study, we do provide evidence that the molecules from ADSCs and FBs are safe for therapeutic development. To help answer these questions, other studies have shown that the secretome from BMSCs induces tumoigenesis in an *in vivo* and *in vitro* mouse model through activation of mTOR (Chaunxia et al, 2018), whereas ADSCs and their conditioned media inhibit cancer growth using *in vivo* and *in vitro* mouse models (Cousin et al, 2009) suggesting that ADSCs are a better cell choice for therapeutic development than are BMSCs in terms of tumorigenesis. ADSCs also present numerous other benefits compared to BMSCs for a variety of safety and efficacy issues (Maguire, 2019). However, more data are needed to understand the long term consequences of the S2RM on various health parameters, and whether injection and IV administration will present a like safety profile as observed in the current studies. In the development of stem cell therapeutics, Goldring et al (2011) have asked the question of whether we are setting a higher bar for the clinical implementation of stem cell-derived therapeutics than we currently apply for other types of cellular therapy. The authors argue there is a danger that if perfection is a prerequisite for beginning stem cell therapeutics, then we will never begin. As Maguire (2019b) has argued, if the current hype about stem cell therapies continues, where unapproved therapies without a knowledge of risk or reward are burgeoning (Turner and Knoepfler, 2016), without a knowledge of the procedure’s risks, then a proper evaluation of the risk versus reward ratio cannot be made for that particular stem cell therapy. Regardless of where the bar is set for other therapies, where medical procedures, like most drugs, have unknown or hidden long term consequences to health, the risks must be evaluated as best as possible so informed risk versus reward decisions can be made. Often, when considering marketed drugs, not until Phase IV, post market approval are the long term consequences of a drug discovered. This is exemplified by the many drugs pulled from market or with safety issues three to four years after their approval (e.g. ProCon, 2014; Downing et al, 2017). Even more unfortunate, the problem is worse with medical procedures (Kumar and Nash, 2011). Such is the case with approved stem cell transplants. Many case studies have reported the approved stem cell transplants to be associated with the later development of cancer (Cooley et al, 2000), and unapproved stem cell transplant procedures are notorious for adverse side-effects, including development of cancer (Diouhy et al, 2014). The effects of approved bone marrow stem cell transplant in cancer relapse are not well understood, but are thought to involve epigenetic factors in the stem cells used for the transplant (Christopher et al, 2018). In addition, bone marrow stem cell transplants may cause aging of the tissue as measured in T-cells using a p16 biomarker (Wood et al, 2016), indicating the increased level of cellular senescence in the surrounding tissue, a factor in the increased probability of tumorigenesis (<u>Schosserer</u> et al, 2017).

While stem cell therapy is in a period of rapid advancement, the science of stem cell safety assessment must also evolve, not to hinder progress, rather to support, guide, and expedite the progress of patient treatment using stem cell-based technologies. The development of a rich safety data base is necessary to ensure that we can proceed with appropriate safeguards in place and allow that stem cell-based therapeutic approaches develop in a way that benefits society overall by using well supported, data driven risk versus reward analysis. These data presented here are one such needed set of safety data to evaluate the patient risk versus benefit ratio for stem cell therapeutics in general, and specifically for our combination of stem cell released molecules from adipose mesenchymal stem cells and fibroblasts.

## Acknowledgements

None

## Author Contributions

Dr. Greg Maguire wrote this paper and helped design and perform the experiments. Peter Friedman helped design and perform the experiments.

## Ethical Approval

The study design and animal usage were reviewed and approved by the CARE Research Institutional Animal Care and Use Committee (IACUC) for compliance with regulations prior to study initiation (IACUC #1730). Animal welfare for this study was in compliance with the U.S. Department of Agriculture’s (USDA) Animal Welfare Act (9 CFR Parts 1, 2, and 3), the Guide for the Care and Use of Laboratory Animals,[1] and CARE Research SOPs.

